# *animalcules*: Interactive Microbiome Analytics and Visualization in R

**DOI:** 10.1101/2020.05.29.123760

**Authors:** Yue Zhao, Anthony Federico, Tyler Faits, Solaiappan Manimaran, Stefano Monti, W. Evan Johnson

## Abstract

**Background:** Microbial communities that live in and on the human body play a vital role in health and disease. Recent advances in sequencing technologies have enabled the study of microbial communities at unprecedented resolution. However, these advances in data generation have presented novel challenges to researchers attempting to analyze and visualize these data.

**Results:** To address some of these challenges, we have developed *animalcules*, an easy-to-use interactive microbiome analysis toolkit for 16S rRNA sequencing data, shotgun DNA metagenomics data, and RNA-based metatranscriptomics profiling data. This toolkit combines novel and existing analytics, visualization methods, and machine learning models. For example, traditional microbiome analyses such as alpha/beta diversity and differential abundance analysis are enhanced in the toolkit, while new methods such as biomarker identification are introduced. Powerful interactive and dynamic figures generated by *animalcules* enable users to understand their data and discover new insights. *animalcules* can be used as a standalone command-line R package or users can explore their data with the accompanying interactive R Shiny interface.

**Conclusions:** We present *animalcules*, an R package for interactive microbiome analysis through either an interactive interface facilitated by R Shiny or various command-line functions. It is the first microbiome analysis toolkit that supports the analysis of all 16S rRNA, DNA-based shotgun metagenomics, and RNA-sequencing based metatranscriptomics datasets. *animalcules* can be freely downloaded from GitHub at https://github.com/compbiomed/animalcules or installed through Bioconductor at https://www.bioconductor.org/packages/release/bioc/html/animalcules.html.

## Background

The complex role of the gut microbiota in shaping human health and disease has been intensely investigated and explored in recent years, largely due to the availability of culture-independent molecular-based high-throughput sequencing technologies. It is estimated that every human host coexists with an average of 500-1000 different bacterial species [1–3] and research has discovered that the microbiome is associated with host lifestyle and diet [4, 5] as well as many diseases such as obesity, type 2 diabetes [6] and cancer [7]. Burgeoning sequencing technology brings not only more data and capacity for microbiome research, but also new challenges for data analytics and interpretation. Improved tools and methods for microbiome data analytics will enhance our ability to understand the roles of microbes in diverse environments, particularly understanding how they interact with each other as well as their human hosts.

Current microbiome analysis typically consists of two important components: upstream community profiling (e.g. what is the abundance of all microbes in each sample?) and downstream high-level analysis (e.g. alpha/beta diversity analysis, differential abundance analysis) [8]. In recent years, evolving data analytics, visualization, and machine learning methods have been gradually applied to the development of many software tools and web servers for microbiome data analysis covering these two components. [9–13]. However, new techniques and sequencing technologies have steepened the learning curve for scientific researchers applying new methods for microbiome data analysis and interpretation [14]. Furthermore, existing tools are mostly dedicated to one aspect of analysis and/or are restricted to analyzing one type of microbiome data. For example, while there are many tools and workflows for analyzing 16S rRNA data, there are no existing tools and pipelines tailored for comprehensively addressing the analytical needs of RNA-based metatranscriptomics.

**Table 1** gives a summary of the functions of these tools with respect to the analysis needs of microbiome data. For marker gene-based data such as 16S rRNA, QIIME II [15] and Mothur [16] provide a user interface and a plethora of analytic and visualization tools, but do not provide support for metagenomic and metatranscriptomic data. Vegan [17] provides a wide variety of functions for metagenomic data visualization, but lacks a user-interface, and tools for host and microbial read alignment, differential expression, etc. BioBakery [18] provides a comprehensive suite of tools for most metagenomic analysis needs for microbial communities, but relies on a small set of markers to identify species, and does not address host or microbial expression. Phyloseq [19] has a Shiny interface with tools for annotation, visualization, and diversity analysis, but does not provide abundance analysis, and is no longer actively maintained by its developers. None of these methods are comprehensive or specifically address the needs for multiple types of 16S rRNA, metagenomic or metatranscriptomic data. Therefore, there are no existing toolkits that contain a complete workflow for microbiome data analysis and interpretation (with or without a graphical user interface).

**Table 1.** Comparison of *animalcules* and other popular microbiome analysis tools.

|  | Biobakery | Vegan | mothur | qiime2 | phyloseq | animalcules |
| --- | --- | --- | --- | --- | --- | --- |
| Filtering and Data Summary | ✓ |  | ✓ | ✓ | ✓ | ✓ |
| Interactive Visualization |  |  |  | ✓ |  | ✓ |
| Dimension Reduction |  |  | ✓ | ✓ | ✓ | ✓ |
| Differential Abundance Analysis | ✓ |  | ✓ | ✓ | ✓ | ✓ |
| Diversity Analysis |  | ✓ | ✓ | ✓ | ✓ | ✓ |
| Support for 16S rRNA Data | ✓ | ✓ | ✓ | ✓ | ✓ | ✓ |
| Support for total RNA-seq Data | ✓ | ✓ |  |  |  | ✓ |
| Biomarker Identification |  |  |  | ✓ |  | ✓ |
| Interface and Command-Line |  |  |  | ✓ |  | ✓ |
| Language/Platform | Web | R | R | Python | R | R |

Here we present *animalcules*, an interactive analysis and visualization toolkit for microbiome data. *animalcules* supports the importing of microbiome profiles in multiple formats such as a species count table, an organizational taxonomic unit (OTU) counts table, or Biological Observation Matrix (BIOM) format [20]. These formats could be generated from common microbiome data sources and analytical tools including 16S rRNA, metagenomics, and metatranscriptomic data. Once data is uploaded, *animalcules* provides a useful data summary and filtering function where users can view and filter their dataset using sample metadata, microbial prevalence or relative abundance. Filtering the data in this way can significantly reduce the time spent performing preprocessing and downstream analysis tasks. For data visualizations, such as relative abundance bar charts and 3D dimension reduction plots (PCA/PCoA/tSNE/UMAP), *animalcules* supports interactive operations where users can check the sample/microbe information on each data point and adjust the figure format as needed, which is helpful for recognizing elements or data patterns when the sample size or number of microbes is large. Aside from common diversity analysis, differential abundance analysis, and dimension reduction, *animalcules* supports biomarker identification by training a logistic regression or random forest model with cross-validated biomarker performance evaluation. *Animalcules* provides a graphical user interface (GUI) through R/Shiny, which can be used even by users without prior programming knowledge, while experienced programmers can choose the command-line based R package or a combination of both.

## Implementation

### Data Structures and Software Design

All data handling tasks and functions in *animalcules* are based upon and work with the MultiAssayExperiment (MAE) data structure [21]. The MAE class is a standard data structure for multi-omics experiments with efficient data retrieval and manipulation methods that support the linkage of samples across multiple assays. The MAE object has three key components: colData (contains subject or cell line level metadata), ExperimentList (stores data for one or more assays), and sampleMap (relates experiments and samples). In *animalcules*, three tables (sample metadata table, microbe count table, and taxonomy table) as well as the mapping relationship between them are stored in the MAE class. It ensures correct alignment of assays and subjects, and provides coordinated subsetting of samples and features. Additionally, it is easy to convert to or from a MAE object from the SummarizedExperiment class, which has been applied in many Bioconductor packages, enabling smooth interaction between other tools [21]. One important advantage of applying the MAE class in the microbiome research field is its extensible design supporting many multi-omics layers of data. Multi-omics is becoming a trend in the field, e.g., studying host-microbe interactions by combining host gene expression data and microbial abundance data. *animalcules* is the first software tool for microbiome analysis to integrate the MAE object and takes advantage of its unique properties by allowing the user to store microbial data, host transcriptomics, metabolomics, as well as taxonomy information within the same object (currently *animalcules* only supports microbiome analysis, but the MAE structure enables future development that can address these data types. Additionally, the MAE enables integration with other tools that do manage these data types, e.g. host transcriptomics). The MAE object can also store processed versions of various assays (e.g. dimension-reduced data) which allows for efficient manipulation and analysis downstream. This approach advances standard microbiome analysis and data sharing by efficiently integrating the various multi-omics datasets required.

Lastly, because all of the data is integrated within a single R object, users can serialize the data to a single file which can be used for further analysis or share with other researchers. For example, after processing and analyzing their data through the Shiny application, users can export their datasets in the form of a serialized MAE object file, which can be later uploaded to Shiny or imported in R for further exploration through the *animalcules* command line functions or other methods. Integrating the MAE object brings efficiency, scalability, and reproducibility to microbiome analysis through *animalcules*.

### Installation and Usage

*animalcules* requires R >= 4.0.0 and can be installed through Github or Bioconductor. After loading the *animalcules* library in R, users can choose between launching the R Shiny GUI (via the run_animalcules() function), or using the available command-line functions directly. In the GUI, users can choose from the following tabs: Upload (select an example dataset, upload a new dataset, or load a previously uploaded dataset), Summary and Filter (understand the data distribution and filter the data by microbial features or sample phenotypes), Abundance (relative abundance bar charts, heatmaps, and individual microbes boxplots), Diversity (statistical tests and boxplots for alpha diversity and beta diversity), Dimension Reduction (PCA, PCoA, tSNE, and UMAP), Differential Abundance (microbial differential abundance between sample groups), and Biomarker (identify predictive microbial biomarkers). Common R functions in the package are summarized in **Table 2**. A detailed tutorial on how to use the command-line version of *animalcules* for microbiome data analysis can be found at https://compbiomed.github.io/animalcules-docs/articles/animalcules.html.

**Table 2.**
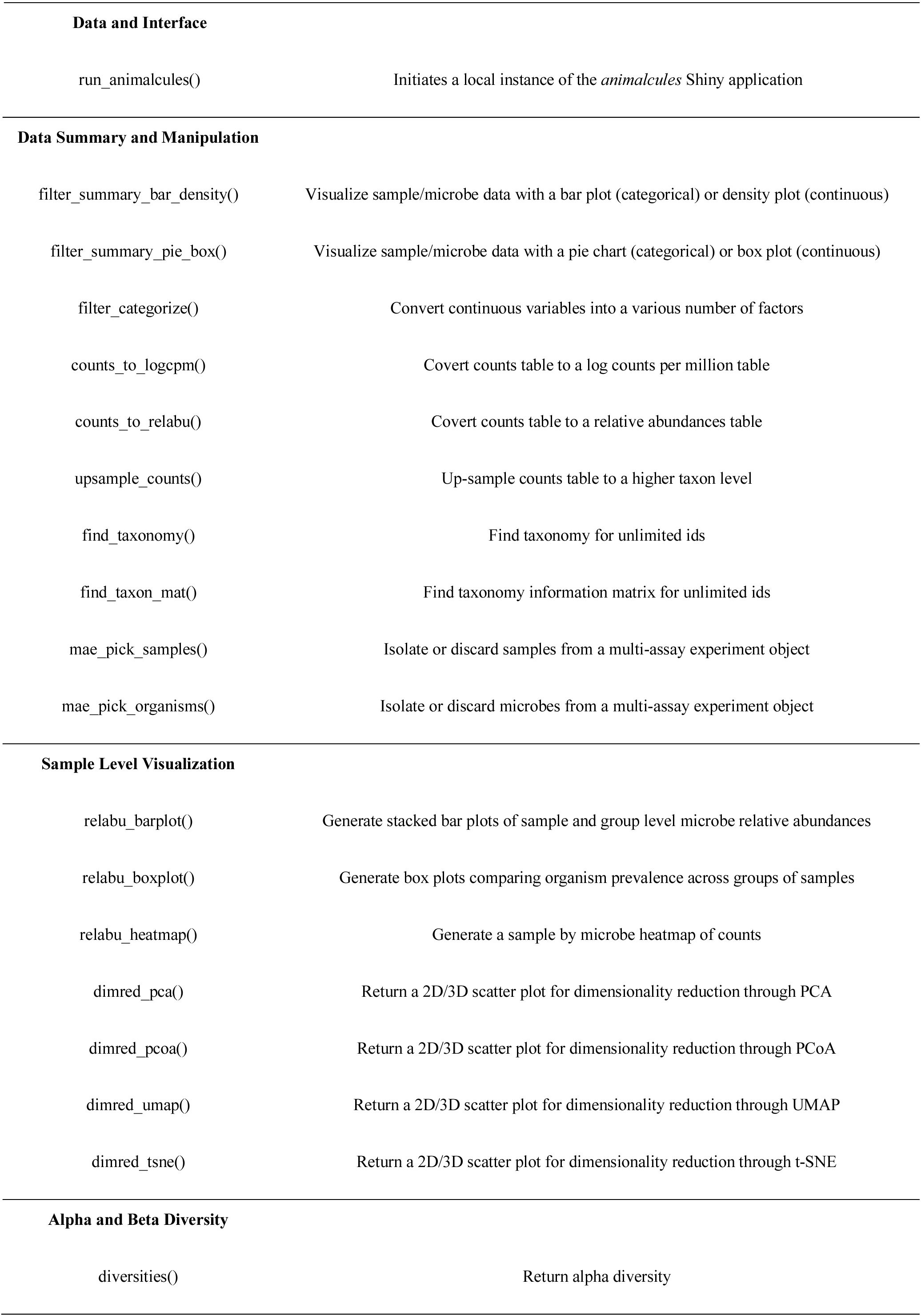
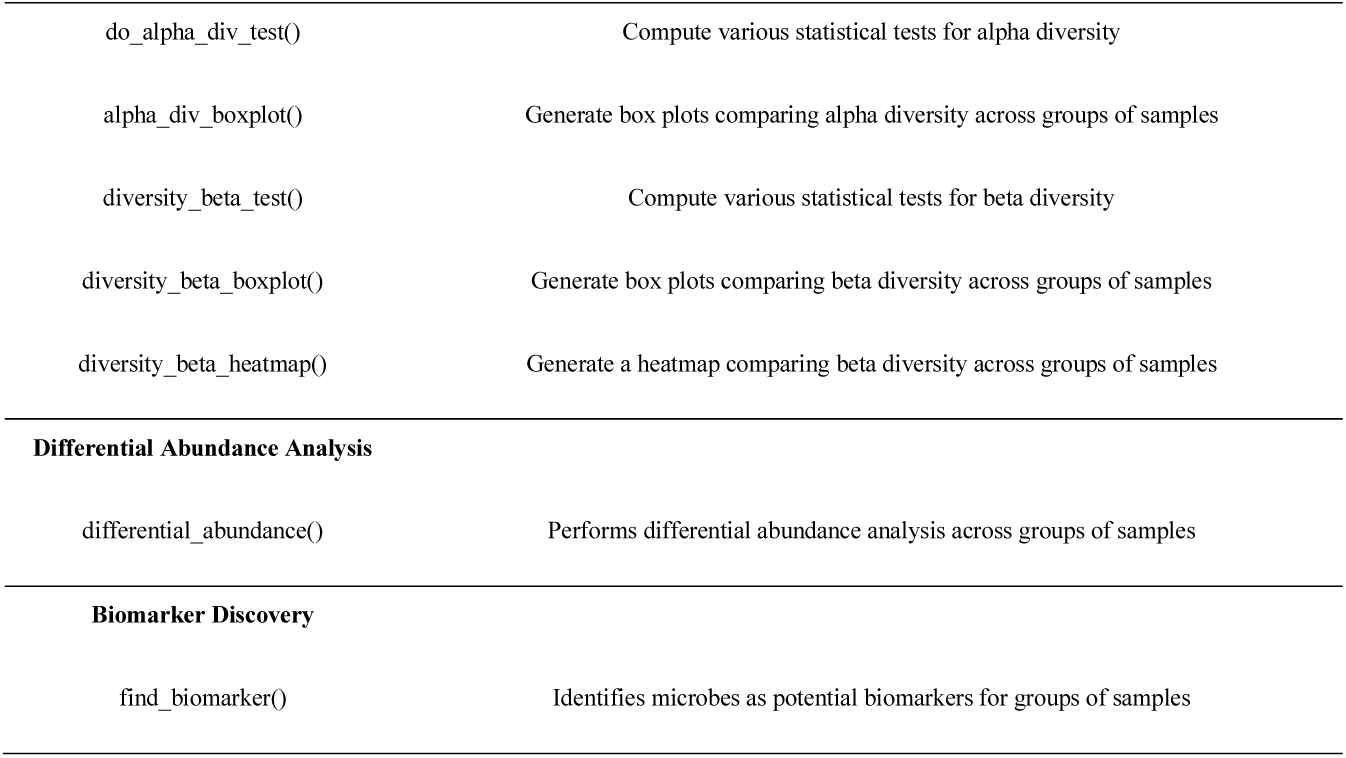
Table of exported functions and their descriptions available through the *animalcules* R package.

Data and Interface
|  |  |
| --- | --- |
| `run_animalcules()` | Initiates a local instance of the *animalcules* Shiny application |
| Data Summary and Manipulation |  |
| `filter_summary_bar_density()` | Visualize sample/microbe data with a bar plot (categorical) or density plot (continuous) |
| `filter_summary_pie_box()` | Visualize sample/microbe data with a pie chart (categorical) or box plot (continuous) |
| `filter_categorize()` | Convert continuous variables into a various number of factors |
| `counts_to_logcpm()` | Covert counts table to a log counts per million table |
| `counts_to_relabu()` | Covert counts table to a relative abundances table |
| `upsample_counts()` | Up-sample counts table to a higher taxon level |
| `find_taxonomy()` | Find taxonomy for unlimited ids |
| `find_taxon_mat()` | Find taxonomy information matrix for unlimited ids |
| `mae_pick_samples()` | Isolate or discard samples from a multi-assay experiment object |
| `mae_pick_organisms()` | Isolate or discard microbes from a multi-assay experiment object |
| Sample Level Visualization |  |
| `relabu_barplot()` | Generate stacked bar plots of sample and group level microbe relative abundances |
| `relabu_boxplot()` | Generate box plots comparing organism prevalence across groups of samples |
| `relabu_heatmap()` | Generate a sample by microbe heatmap of counts |
| `dimred_pca()` | Return a 2D/3D scatter plot for dimensionality reduction through PCA |
| `dimred_pcoa()` | Return a 2D/3D scatter plot for dimensionality reduction through PCoA |
| `dimred_umap()` | Return a 2D/3D scatter plot for dimensionality reduction through UMAP |
| `dimred_tsne()` | Return a 2D/3D scatter plot for dimensionality reduction through t-SNE |
| Alpha and Beta Diversity |  |
| `diversities()` | Return alpha diversity |
| `do_alpha_div_test()` | Compute various statistical tests for alpha diversity |
| `alpha_div_boxplot()` | Generate box plots comparing alpha diversity across groups of samples |
| `diversity_beta_test()` | Compute various statistical tests for beta diversity |
| `diversity_beta_boxplot()` | Generate box plots comparing beta diversity across groups of samples |
| `diversity_beta_heatmap()` | Generate a heatmap comparing beta diversity across groups of samples |
| Differential Abundance Analysis |  |
| `differential_abundance()` | Performs differential abundance analysis across groups of samples |
| Biomarker Discovery |  |
| `find_biomarker()` | Identifies microbes as potential biomarkers for groups of samples |

### Data Upload and Output

*animalcules* offers multiple options for importing data into the GUI or working with the MAE object for command line analysis. These include simple tab-delimited OTU or count matrices, typically generated by other tools such as QIIME II [15] or PathoScope [22], or using a MAE object available in the user’s session or in a file from a previous session of *animalcules.* Regardless of how the data is imported, the assay/OTU data will be available in the “Assay Viewer” section of the Upload tab.

Six of the data importing options are described below:

1. *Count Table or OTU File (without taxonomy):* This is the simplest option that enables the upload of an OTU or count table that has genomes/OTUs in the rows and samples in the columns. All functions and tools can be used for filtering, visualization, analysis of the data, except the individual microbiomes or OTUs cannot be aggregated at different levels.
2. *Count Table or OTU File (with taxonomy):* This option provides an extension of the previous but allows for associating the OTUs with taxonomy information and the aggregation of microbes at different levels (e.g. species, genus, phylum, etc.). This information can be provided as a separate table, with a row for each OTU in the table. In addition, users can provide NCBI taxonomy IDs or NCBI accession numbers [23] and *animalcules* will automatically generate the taxonomy table using the tools available in the *taxize* R package [24]. The taxonomy table will be stored as a separate assay in the MAE object, but will be linked to the rows of the OTU table through internal functions. The taxonomy table will be available in the “Assay Viewer” section in the Upload tab.
3. *animalcules Object File:* Users can also directly upload a MAE object into the toolkit or workflow. A MAE object could be generated from a previous *animalcules* session (stored as an .rds file), converted from the output of any pre-processing pipeline, or generated from some other source. This option allows for the efficient storage and re-upload of data from a previous session, or enables the interaction between the command-line version and the GUI version of animalcules. For example, users can conduct part of the analysis in the GUI, save the results, and continue their analysis using command-line tools (inside and outside of *animalcules*), and then re-upload the data to the GUI for further analysis or visualization. This feature enables compatibility and interactivity that is not available in other microbiome GUI or command-line toolkits.
4. *Pathoscope Output Files: animalcules* enables the direct upload of files generated from the *PathoScope* pipeline [22]. These files are generally single tab-delimited tables for each sample in the dataset, and contain NCBI taxonomy IDs for individual microbes. *animalcules* combines and converts these files into a MAE object, and uses the *taxize* package to generate the taxonomy table.
5. *BIOM Format File:* The standard BIological Observation Matrix (BIOM) format is a commonly used format for representing samples by observation contingency tables [20]. The BIOM format is commonly used by QIIME II pipeline tools. We used the *biomformat* R package [25] for uploading a BIOM file into *animalcules* as well as outputting a BIOM file from *animalcules*. This enables interactivity between *animalcules* and other microbiome analysis tools such as QIIME II.
6. *Example Data*: In *animalcules*, we have three pre-defined example datasets, including a simulated dataset, a Tuberculosis 16S rRNA profiling dataset, and an Asthma metatranscriptomic dataset. These example datasets allow users to try all the features and functions in animalcules before users upload their own data, making it easy to learn how to use *animalcules* and understand what analyses they can perform.

### Data Filtering and Summary

The *animalcules* Shiny interface provides summary statistics to help users efficiently and effectively assess data quality and filter low-quality microbes and samples. Users can visualize the total number of reads for each organism through a scatter and density plot and filter organisms based on average read number, relative abundance, or prevalence. Additionally, users can visualize sample covariates through a pie and bar plot for categorical covariates or a scatter and density plot for continuous covariates **(Figure 1)**. Samples can be filtered based on one or more covariates. Finally, users have the option to discard specific samples and/or organisms. As samples and organisms are removed through any of the filtering methods, summary statistics and plots are automatically refreshed to display any changes that may occur. If changes have been made, users may download the modified data for later use. Visualizations of sample and microbe data before and after filtering are generated with animalcules::filter_summary_bar_density() and animalcules::filter_summary_pie_box() functions. For users who wish to inspect their data before or after filtering, *animalcules* enables users to view and download five types of assays generated including a count table, relative abundance table, logCPM table, taxonomy table and annotation table. In addition, these tables can also be accessed directly from the MAE object through standard R command line tools.

**Figure 1.**
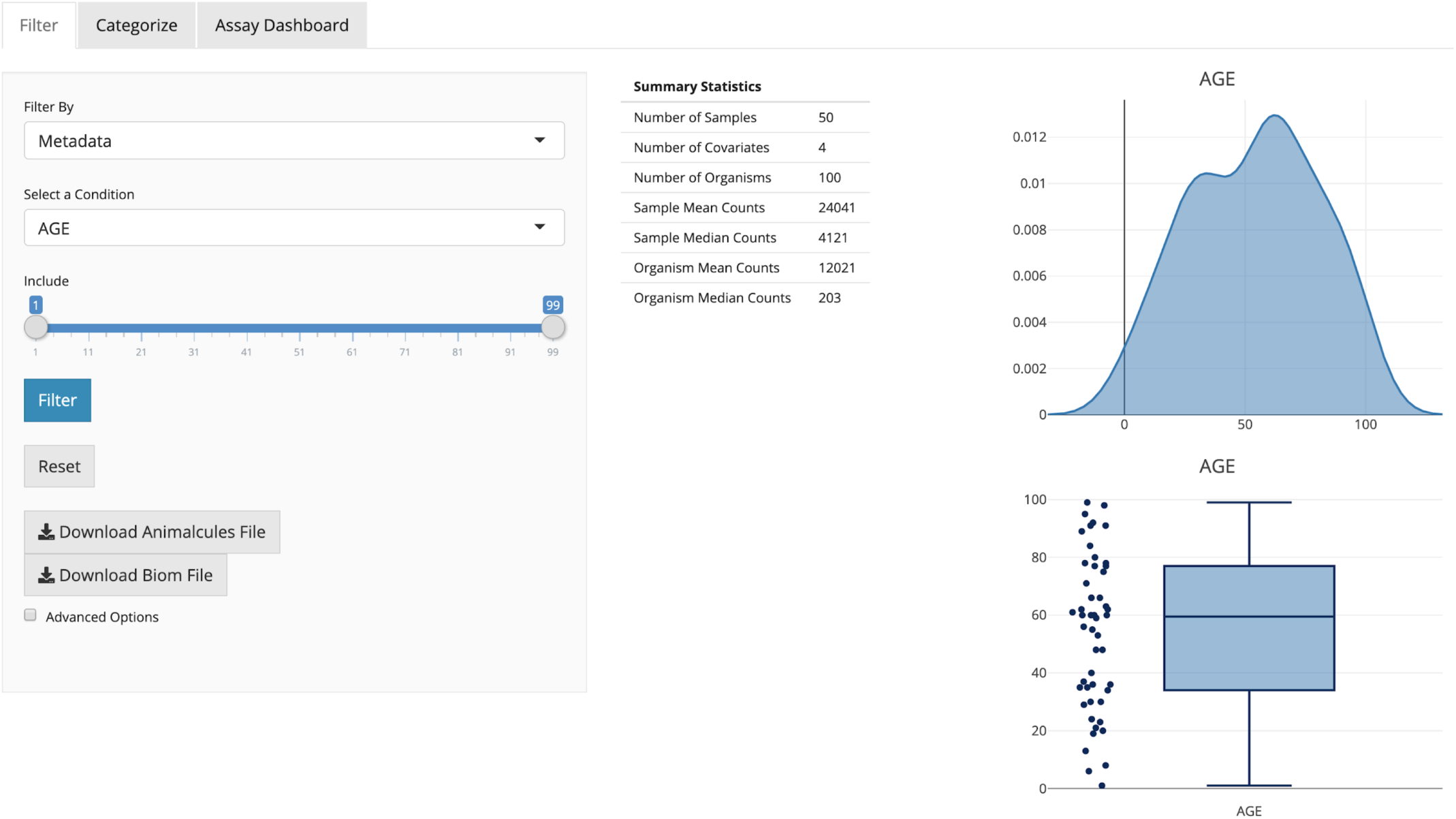
*animalcules* Data Filtering and Summary tab. In the right panel, a table of data summary metrics, a scatter/boxplot, and a density plot are displayed for continuous variables. For categorical variables, *animalcules* will automatically identify and show the pie and bar plots instead.

### Data Visualization

A typical analysis involves visualization of microbe abundances across samples or groups of samples. *animalcules* implements three common types of visualization plots including stacked bar plots, heatmaps, and box plots. The stacked bar plots, generated with animalcules::relabu_barplot() are used to visualize the relative abundance of microbes at a given taxonomic level in each sample, represented as a single bar **(Figure 2)**. Bars can be color-labeled by one or more sample attributes and samples can also be aggregated by these attributes via summing microbe abundances within groups. This is an efficient way for researchers to identify sample- or group-level patterns at various taxonomic levels. Users also have the option to sort the bars by sample attributes or by the abundance of one or more organisms. There is also a convenient method for isolating or removing samples. With this tool, users can quickly scan through different combinations of sample attributes and taxon levels for differential abundance in one or more groups, outliers in terms of community profile, as well as sample clusters not represented by known attributes.

**Figure 2.**
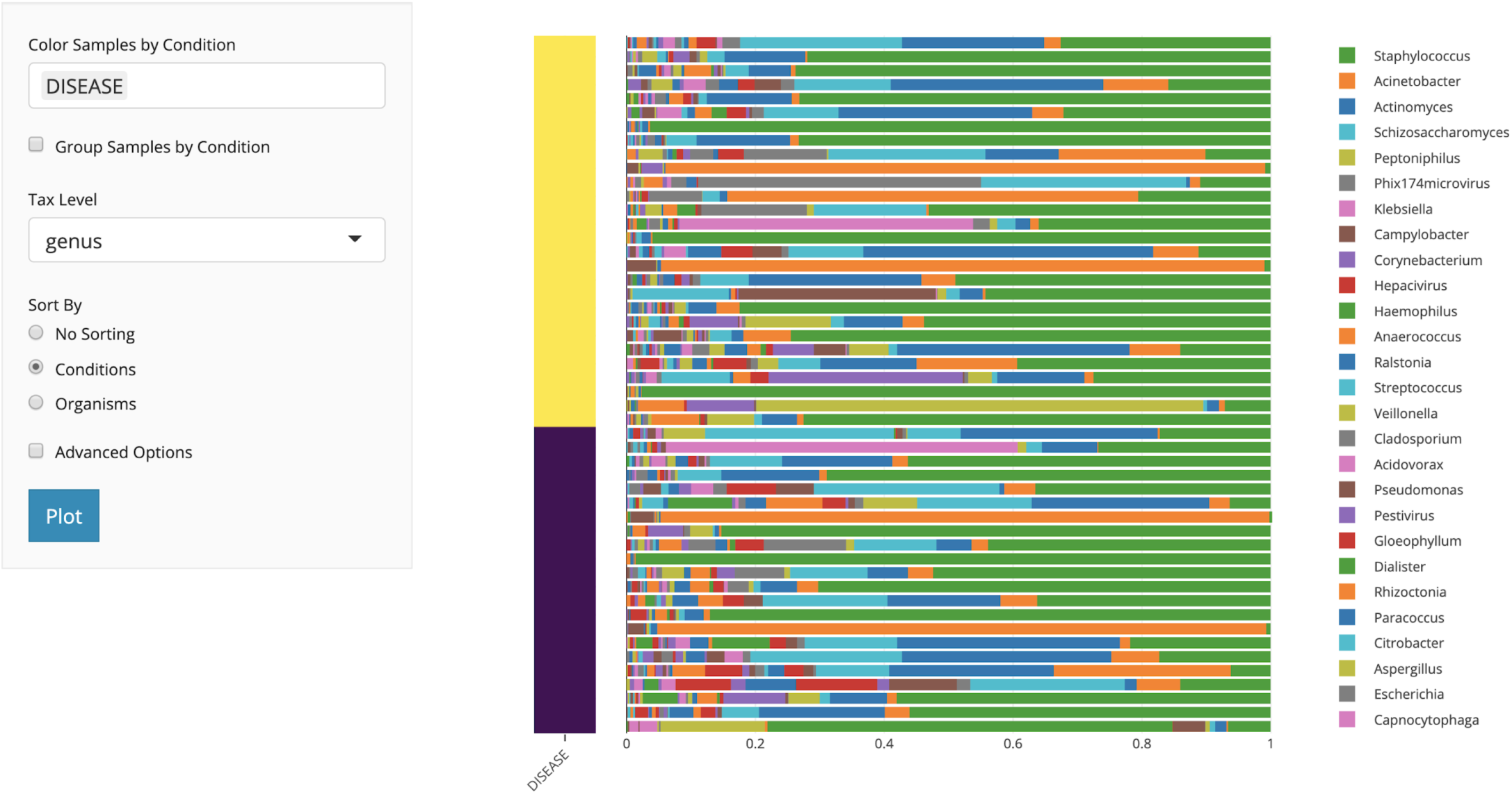
*animalcules* Abundance Tab. In the subtab panel, users can select between a bar plot, heatmap, or box plot. In the bar plot setting, in the left panel, users can select the color by variable, taxonomy level, and sort by option. In the right panel, *animalcules* will show an interactive plot where users can mouse-hover to check the identity of any color bar shown in the plot.

Alternatively, users can investigate these questions through the heatmap visualization, which represents a sample-by-organisms matrix that can be visualized at different taxonomic levels. Many of the previously mentioned options are also compatible with the heatmap such as color-labeling samples, sorting matrix rows by attributes or organisms and isolating or discarding organisms and samples. After identifying potential differentially abundant microbes, users can use the boxplot visualization to directly compare the abundance of one or more organisms between categorical attributes. Organisms can be chosen from a given taxonomic level and abundance can be represented as either counts, logCPM, or relative abundance. This plot can also be generated in the command line using the animalcules::relabu_heatmap() function.

### Diversity Analysis

Alpha diversity, which describes the richness and evenness of a microbial community, is a vital indicator and measurement in microbiome analysis [26]. *animalcules* provides an interactive box plot comparison of alpha diversity between selected groups of samples. Both taxonomy levels and alpha diversity metrics (e.g. Shannon, Gini Simpson, Inverse Simpson) can be changed and diversity can be calculated at multiple taxonomic levels [27, 28]. Alpha diversity values for each sample can be output into the MAE object or as separate tables or files. Users can also conduct alpha diversity statistical tests including Wilcoxon rank-sum test, T-test and Kruskal-Wallis test [29, 30].The alpha diversity boxplot as well as the statistical tests could be generated in the command line using the animalcules::alpha_div_boxplot() function and animalcules::do_alpha_div_test() function.

On the other hand, one can use distances between each microbial community sample, or so-called beta diversity, as another key metric to consider for each analysis. Users can plot the beta diversity heatmap by selecting different beta diversity dissimilarity metrics including Bray-Curtis [31] or Jaccard index [32]. Users can also conduct beta diversity statistical testing between groups including PERMANOVA [33], Wilcoxon rank-sum test, or Kruskal-Wallis test **(Figure 3)**. The beta diversity comparison boxplot as well as the statistical tests can be generated in the command line using the animalcules::diversity_beta_boxplot() function and animalcules::diversity_beta_test() function.

**Figure 3.**
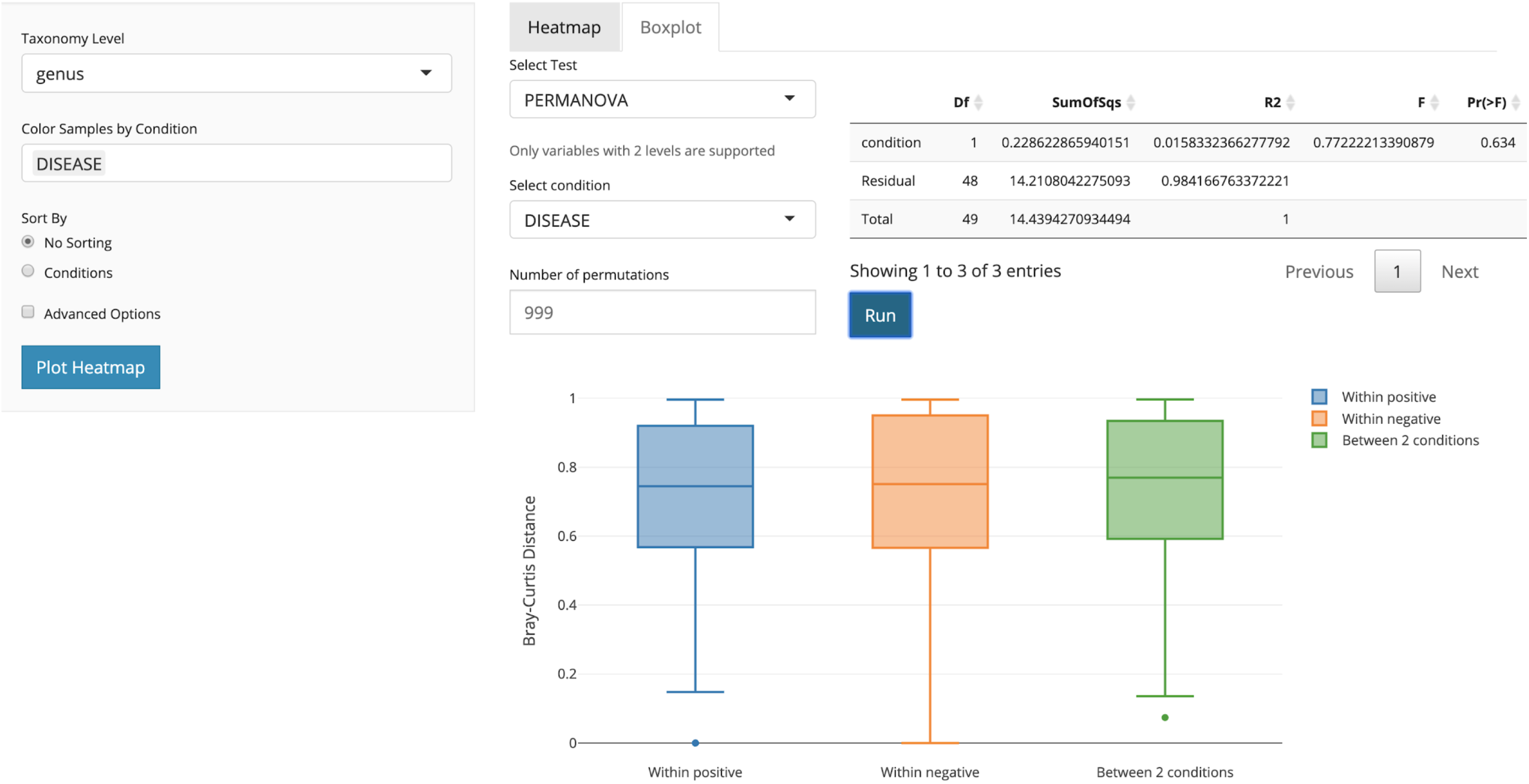
*animalcules* Diversity tab. In the subtab panel, the user could select between alpha diversity analysis and beta diversity analysis. Here in the beta diversity analysis, the right panel controls what statistical test to use, which condition to test on, and show statistical test results in a table as well as a boxplot.

### Dimension Reduction

A crucial step in any data analysis workflow is to visualize and summarize highly variable data in a lower-dimensional space **(Figure 4)**. In *animalcules*, we implement four commonly used dimensionality reduction techniques including Principal Components Analysis (PCA), Principal Coordinates Analysis (PCoA), t-Distributed Stochastic Neighbor Embedding (t-SNE) and Uniform Manifold Approximation and Projection (UMAP) [34–37] Both PCA and PCoA project samples onto a new set of axes whereby a maximum amount of variation is explained by the first, second, and third axes while t-SNE and UMAP are non-linear methods for mapping data to a lower-dimensional embedding. Dimension reduction values for the dataset can be output into the MAE object or as separate tables or files.

**Figure 4.**
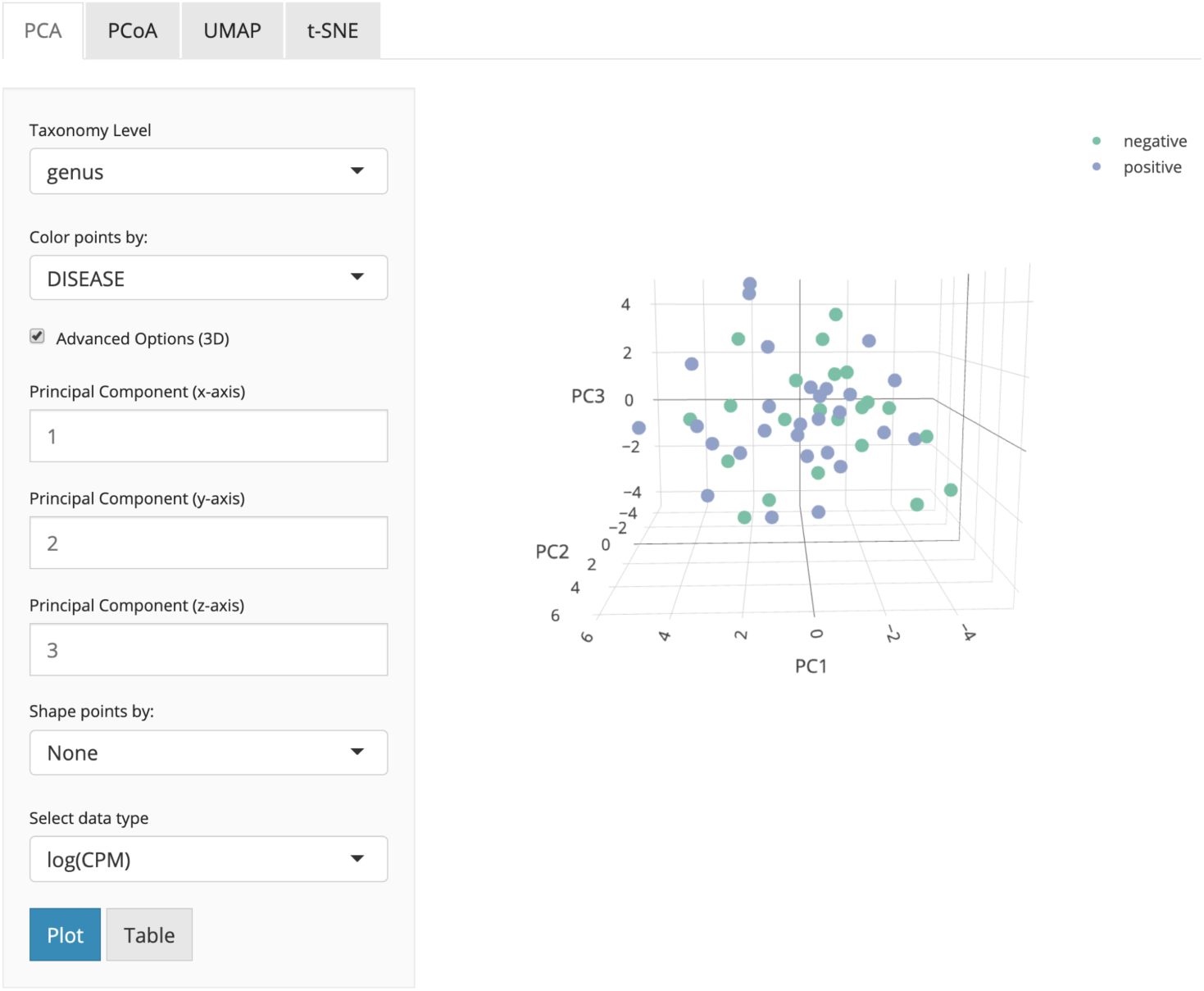
*animalcules* Dimension Reduction tab. In the subtab panel, the user could select between PCA, PCoA, t- SNE, and UMAP. Here in the PCA subtab, the user could choose the taxonomy level, color by variable, and in advanced options, the user could also specify up to three PCs for visualization, shape by variable, and which data type to use.

The original data used in each dimensionality reduction method can be either counts, logCPM, or relative abundance, and can be visualized using a 2D or 3D (if two dimensions of explained variance are inadequate) scatter plot. Data points can be colored by continuous sample attributes and shaped by categorical attributes. With multiple dimensionality reduction techniques and methods for data normalization, users can rapidly visualize the global and local structure of their data, identify clustering patterns across one or more conditions, as well as detect sample outliers. Dimensionality reduction can also be carried out in the command line using the animalcules::dimred_pca() function for PCA, animalcules::dimred_pcoa() function for PCoA, animalcules::dimred_tsne() function for t-SNE, and animalcules::dimred_umap() function for UMAP.

### Differential Abundance Analysis

There are many available tools for differential abundance estimation and inference. For example, GLM (Generalized Linear Model) based methods including DESeq2 [38], edgeR [39], and limma [40] model count based microbiome data or gene expression data by a negative binomial distribution (DESeq2 and edgeR) or using log-counts (per million) and a Gaussian distribution (limma) assumption. Core microbes that have different abundance in different groups could be identified. Here in *animalcules*, we provide a DESeq2-based differential abundance analysis **(Figure 5)**. With the command-line function animalcules::differential_abundance(), which by default uses the “DESeq2” method. Users can choose the target variable, covariate variable, taxonomy level, minimum count cut-off, and an adjusted p-value threshold. The analysis report will output not only the adjusted p-value and log2-fold-change of the microbes, but also the percentage, prevalence, and the group size-adjusted fold change. Besides using DESeq2, in *animalcules* we also support differential abundance analysis with limma, which requires users to specify in the command-line function as: animalcules::differential_abundance(method=‘limma’).

**Figure 5.**
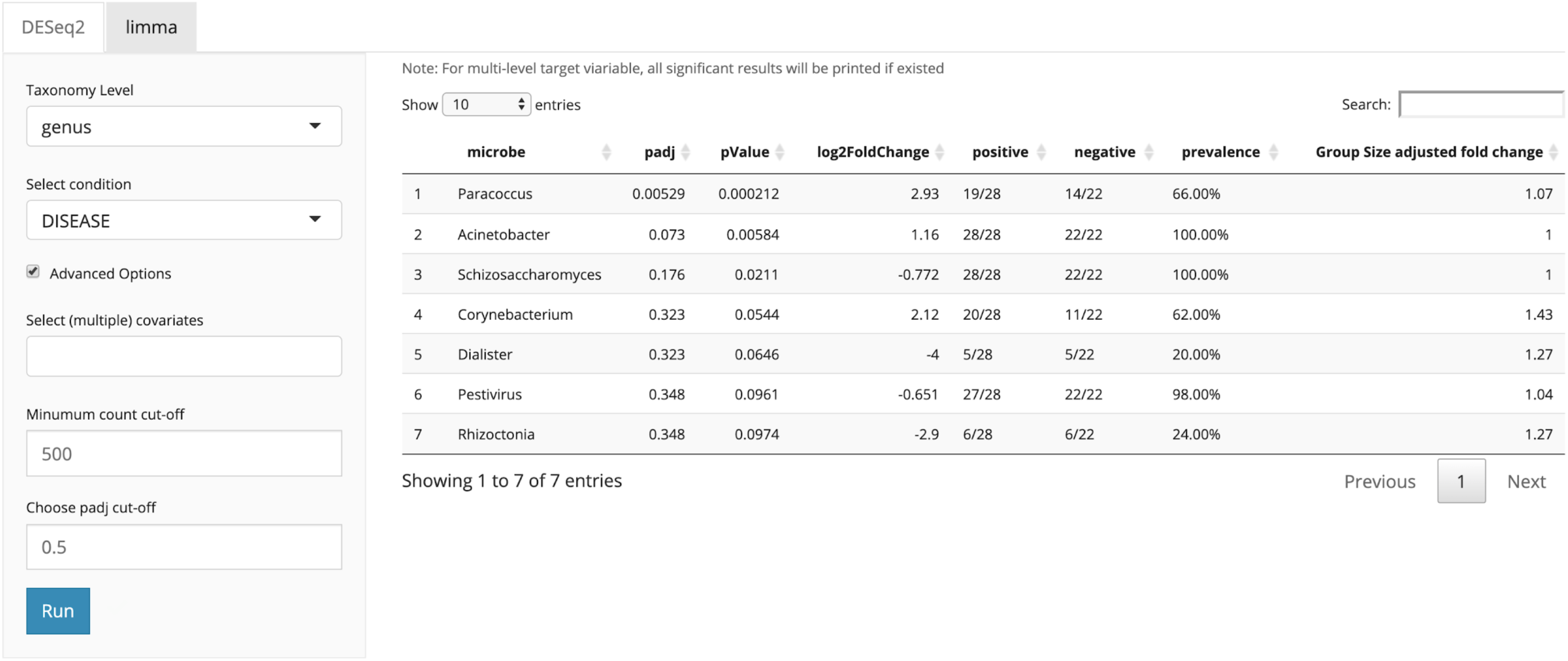
*animalcules* Differential Abundance tab. In the subtab panel, users select between DESeq2 and limma. In the left panel, users specify taxonomy level, target condition, covariate variables, count cut-off, and adjusted p-value threshold. In the right panel, a detailed differential abundance result table is shown.

### Biomarker Identification

One unique feature of *animalcules* is the biomarker identification module. Users can choose either logistic regression [41] or random forest [42] classification model to identify a microbe biomarker. The feature importance score for each microbe will be provided **(Figure 6)**, in addition to AUC values and average cross-validation ROC curves for evaluating biomarker prediction performance. The biomarker identification can also be conducted by the command-line function animalcules::find_biomarker().

**Figure 6.**
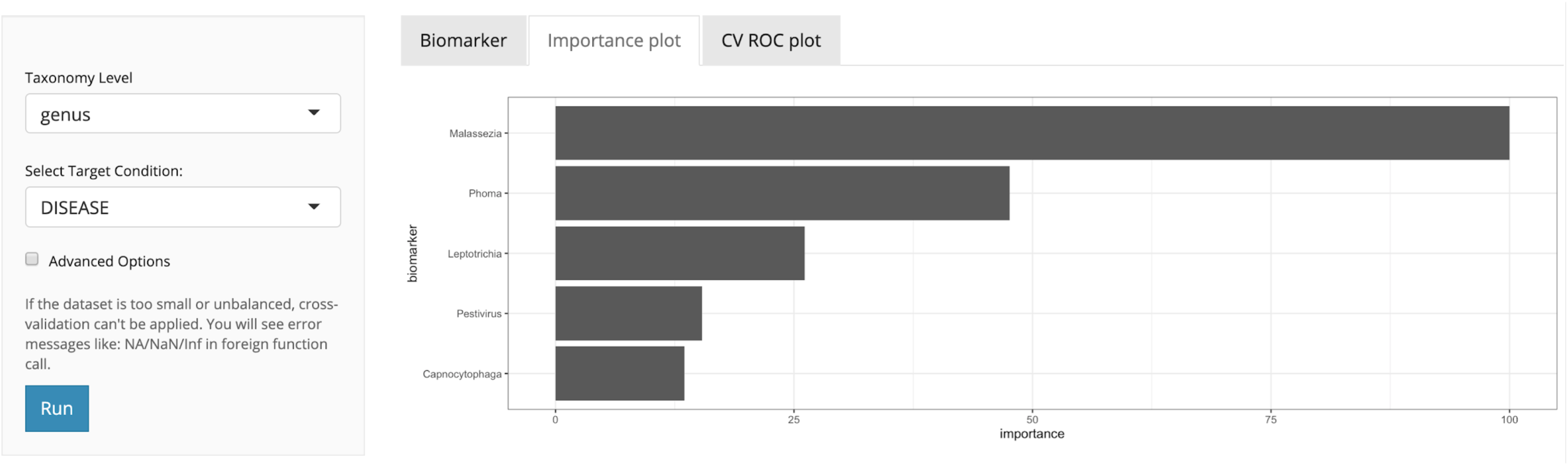
*animalcules* Biomarker tab. In the left panel, users select taxonomy level, and target condition. In the advanced options: number of cross-validations folds, number of cross-validations repeats, biomarker proportion, and classification model. In the right panel, *animalcules* will show the biomarker list, importance plot, and ROC plot.

## Results

To illustrate the utility of *animalcules*, we include two example analyses using the pre-loaded datasets packaged within *animalcules*; the first being an asthma metatranscriptomic dataset, and the second a TB 16S rRNA dataset. For brevity, we do not explore all *animalcules* functions in each analysis, but focus on and expand the relevant analyses for the scientific questions for each example. Both analyses could be reproduced within the *animalcules* Rshiny app by using the corresponding example datasets.

### Example 1: Asthma nasal swabs metatranscriptomic dataset

The asthma metagenomic shotgun RNA sequencing dataset was generated from participants of the AsthMap (Asthma Severity Modifying Polymorphisms) project and originally reported in a research article characterizing asthma-associated microbial communities [43]. It contains 14 total samples of nasal epithelial cells collected from 8 children and adolescents with asthma and 6 healthy controls. The goal of this study was to further understand the relationship between the microbiome and host inflammatory processes in asthmatic children.

To characterize the relationship between microbial communities and asthma, species-level abundances were visualized by plotting the group-wise relative abundance of microbes across asthma and control subjects. This plot can be generated with the animalcules::relabu_barplot() function as well as under the *Abundance* tab of the Shiny application. It is clear that *Moraxella catarrhalis* is overrepresented in asthmatics versus controls, which was a major discovery in the original publication. This microbe - which is known to cause infections in the respiratory system - could serve as a biomarker for early disease detection, severity of disease, or potential for exacerbation. In addition, other dysbiosis to the airway microbiome included differences in other genera such as *Corynebacterium aurimucosum*, which is underrepresented in asthmatics versus controls (**Figure 7**).

**Figure 7.**
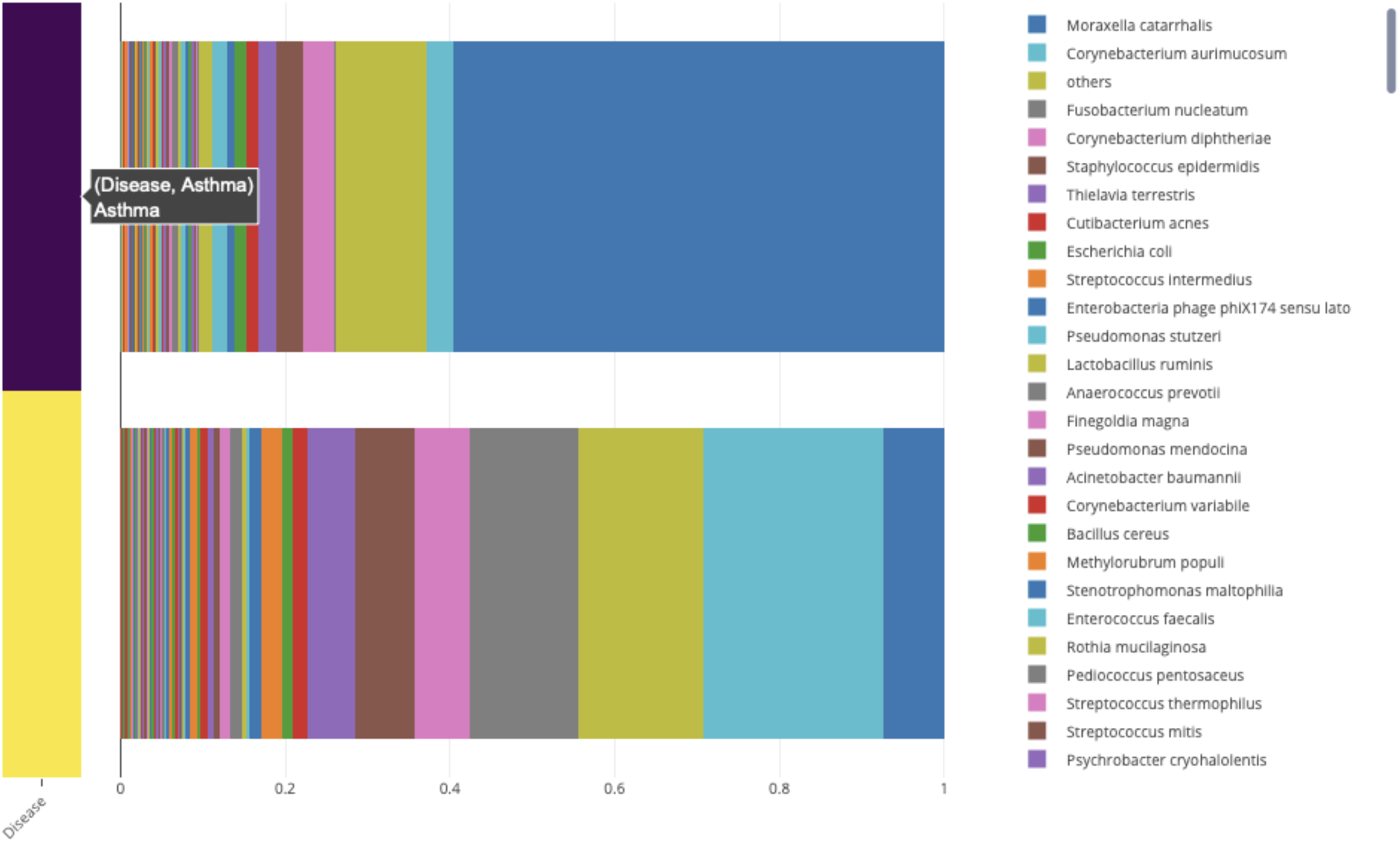
Relative abundance of microbial species bat plot. A stacked bar plot representing the group-wise relative abundance of microbial species in asthmatics (purple) and healthy controls (yellow).

To further investigate the overrepresentation and underrepresentation of *M. catarrhalis* and *C. aurimucosum* respectively in asthmatics versus controls, we use boxplots, generated with the animalcules::boxplot() function, to visualize the relative abundance in each group and to get a better sense of the mean and variance of the distribution across samples. These plots confirm the previous results by showing a drastic difference in abundance (**Figure 8**). Furthermore, we employed DESeq2 to conduct a differential abundance analysis of microbe species for asthmatics versus controls. This analysis shows that *M. catarrhalis* is significantly (q = 1.78e-3) overrepresented (Log_2_FC = 5.9) in asthmatics. It also shows that *C. aurimucosum* is overrepresented (Log_2_FC = 2.66) in controls, however not at a statistically significant level (q = 0.236). This table was generated with the animalcules::differential_abundance() feature.

**Figure 8.**
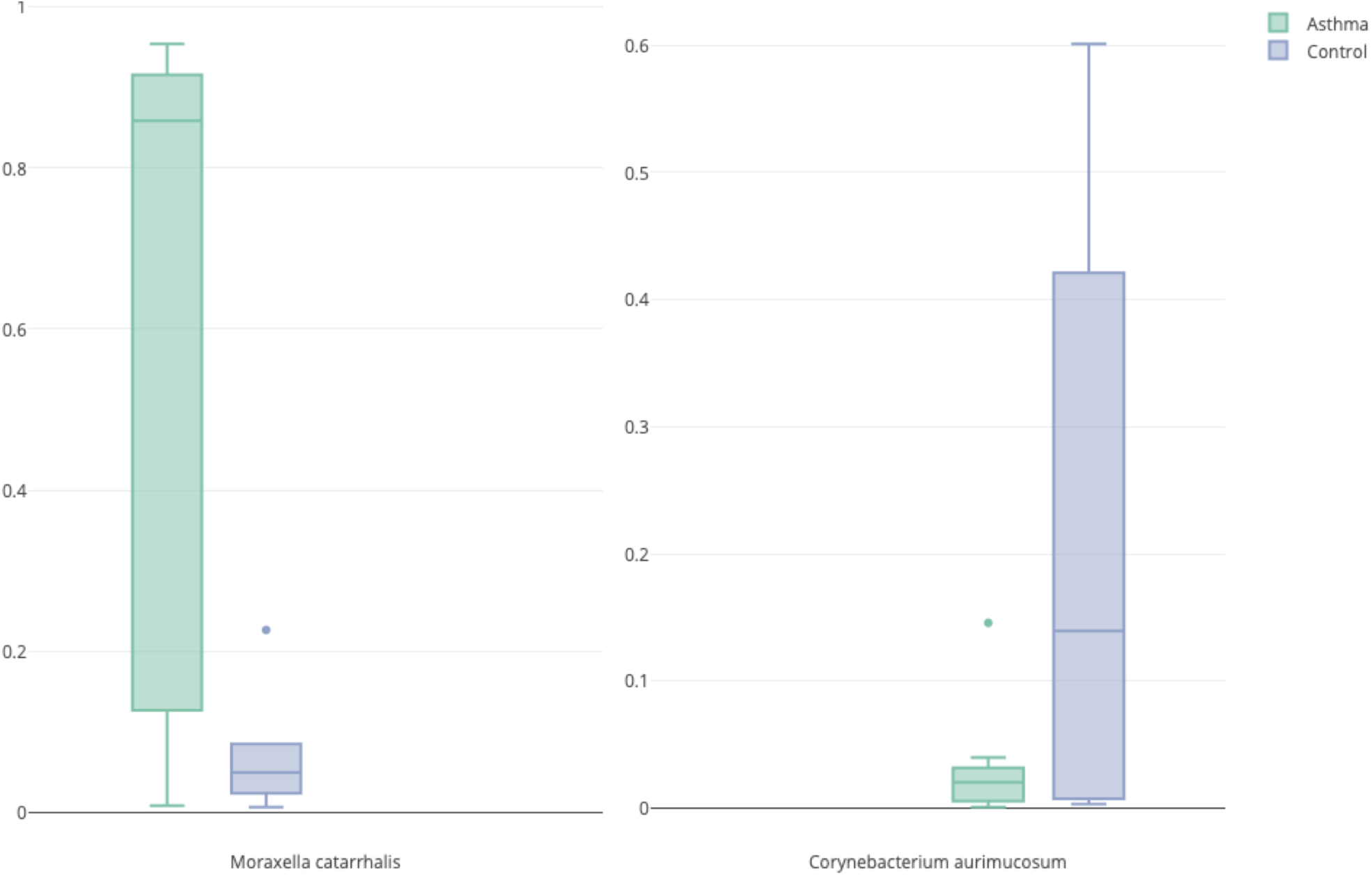
Relative abundance boxplot for differentially abundant species. *Left.* A boxplot of relative abundance of *M. catarrhalis* in asthmatics (green) and healthy controls (blue). *Right.* A boxplot of relative abundance of *C. aurimucosum* in asthmatics (green) and healthy controls (blue).

Through the *animalcules* interface, we were able to rapidly visualize sample- and group-level microbial communities between asthmatic and control samples and test for over- and underrepresented organisms in asthmatics, identifying *M. catarrhalis* and *C. aurimucosum* respectively.

### Example 2: Tuberculosis 16S rRNA profiling dataset

This 16S rRNA TB dataset comes from a pilot TB study containing 12 subjects, 30 respiratory tract samples and 417 species of microbe [44] Among the 12 subjects, there are 6 patients with pulmonary tuberculosis and 6 healthy control individuals. Sample tissue type includes sputum, oropharynx, and nasal respiratory tract. The goal of this study is to learn the microbial community differences in the respiratory tract between healthy and TB patients, and to evaluate the sample/tissue types that were most effective for exploring differences between the microbiome of TB samples vs. controls.

We first conducted an overall assessment of the data, focusing on how the microbial taxonomy affects sample variables such as disease status. We used the barplot function in *animalcules* to visualize the taxonomic profile for each sample, colored by any annotation variable (here we used disease information, where dark blue represents control and yellow represents TB samples). In **Figure 9** we display the genus and phylum level abundances.

**Figure 9.**
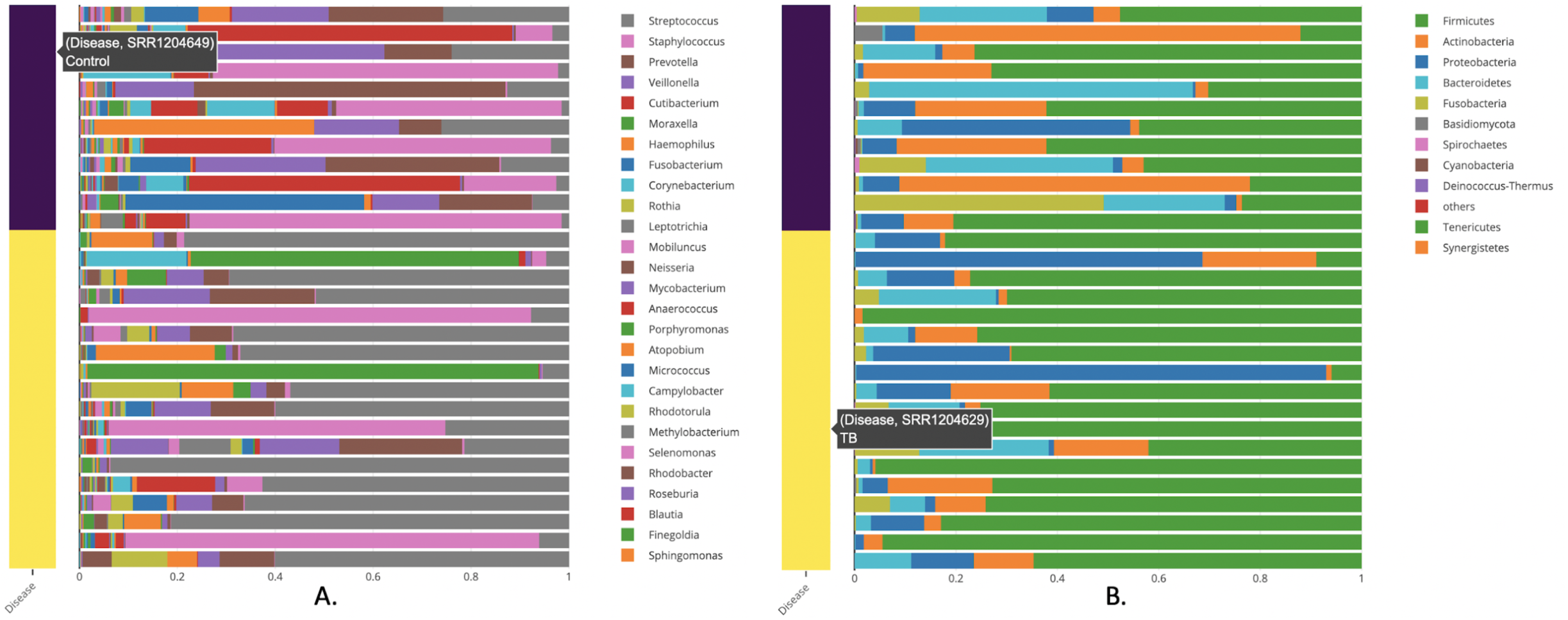
Sample-wise relative abundance bar plot. A stacked bar plot representing the sample-wise relative abundance of microbial species in TB (yellow) and healthy controls (blue). Figure A is the genus level and figure B is the phylum level.

From the taxonomy barplot, we find different patterns that exist in TB vs. control samples. At the genus level (**Figure 9A**), *Streptococcus* appears to have a higher relative abundance in TB samples compared to the control samples. In the phylum level (**Figure 9B**), we found *Firmicutes* to be more abundant in TB samples. Both figures were generated using command-line function animalcules::rebalu_barplot().

To obtain a quantitative understanding of the ecological diversity difference between TB and control samples, we compared the alpha and beta diversity of our samples. For alpha diversity, we compared the Shannon index in TB vs. control samples (see **Figure 10.A**). *animalcules* automatically conducted a non-parametric Wilcoxon rank-sum test and a parametric Welch two-sample T-test on these diversity measures. Here the Wilcoxon rank-sum test gives a p-value of 0.0060, while Welch two-sample T-test gives a p-value 0.0077, thus showing a significant difference in diversity between TB and control groups. From the boxplot, we observe that the alpha diversity is higher in the control group. The alpha diversity boxplot was generated by animalcules::alpha_div_boxplot(), and the statistical test was generated by animalcules::do_alpha_div_test().

**Figure 10.**
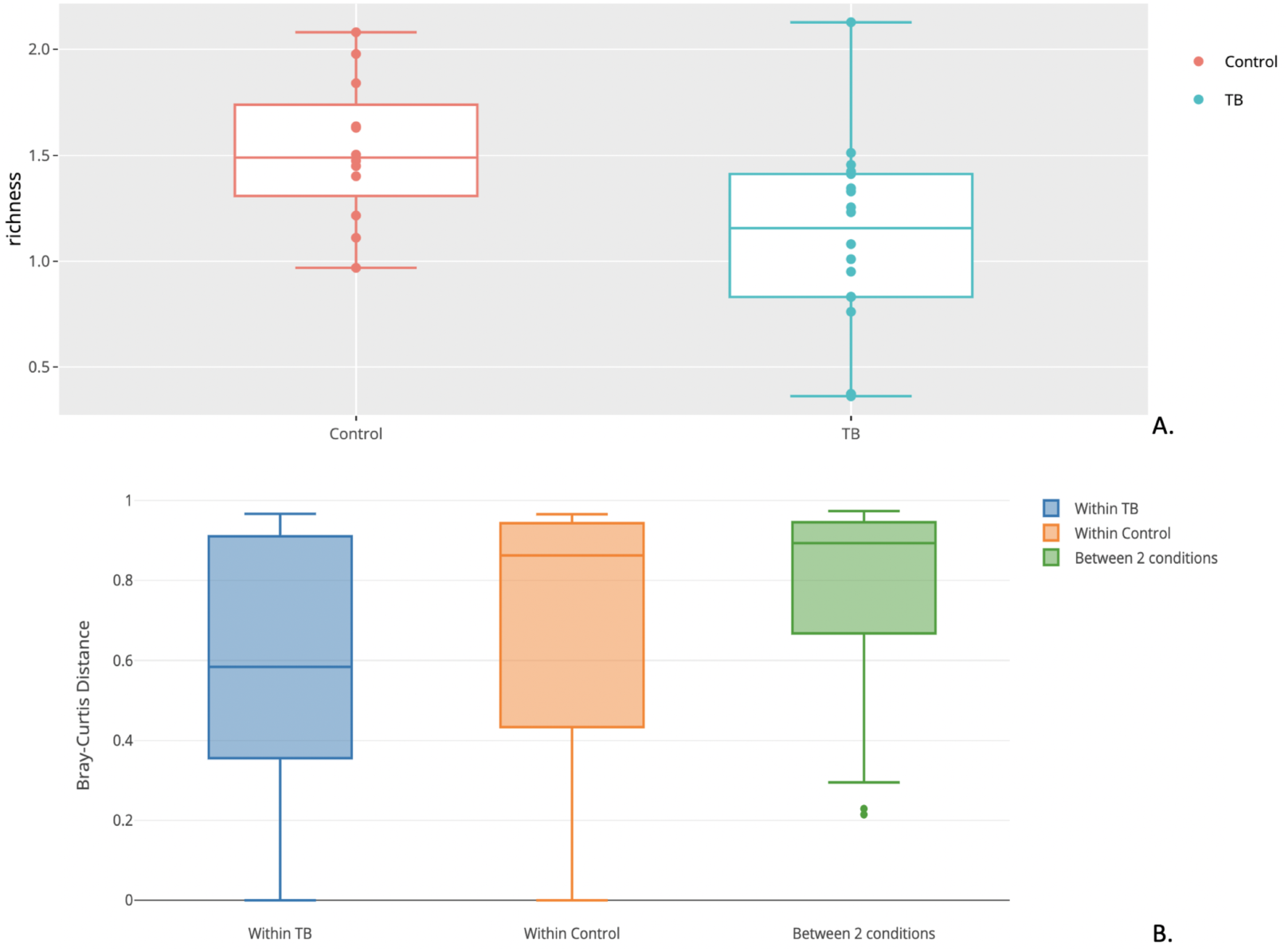
TB example dataset diversity analysis. **A**. Alpha diversity boxplot between control (red) group and TB (blue) group. **B**. Beta diversity boxplot within the TB (blue), within the control(orange), and between TB/control group (green).

As for comparing beta diversity, we plotted the Bray-Curtis distance to compare: within the TB group, within the control group, and between the TB/control (**Figure 10.B**). The average distance between the two groups is higher than two separate within-group distances, meaning both TB samples and control samples are more similar to themselves. Furthermore, we conducted a PERMANOVA test between the two groups, and it shows a significant difference with a p-value of 0.003. The beta diversity comparison boxplot was generated by animalcules::diversity_beta_boxplot(), and the PERMANOVA test was generated by animalcules::diversity_beta_test().

After exploring this TB dataset in terms of relative abundance and diversity analysis, we were certain that there is a significant difference between TB and control groups in the microbiome. Here, with the biomarker function in *animalcules*, we were able to build a microbial biomarker that could help us predict TB status. Using a logistic regression model, 3-fold cross-validation (CV), the number of CV repeats as 3, and top biomarker proportion as 0.05, we identified an 8-genus biomarker for TB classification. Then we tested the biomarker performance by using only the 8-genus biomarker for cross-validation, and the prediction performance ROC is displayed in (**Figure 11)**. We used animalcules::find_biomarker() to identify the biomarker, plot the feature importance score barplot and the ROC curve. Here we have a very high AUC = 0.913, thus providing evidence that the microbiome could serve as a biomarker for TB prediction, and our biomarker has a differentiating power between TB vs. healthy controls. This result suggests that further evaluation of microbial biomarkers for TB is warranted. Previously, people have been using transcriptomic biomarkers for TB diagnosis [45], our new finding of using microbes as diagnosis biomarker can lead to a potential total RNA-seq directed TB disease biomarker that involves both host transcriptome gene expression as well as microbial abundance, which has the potential of higher accuracy for TB diagnosis because it considers both host and microbial side, or even the host-microbe interaction in TB.

**Figure 11.**
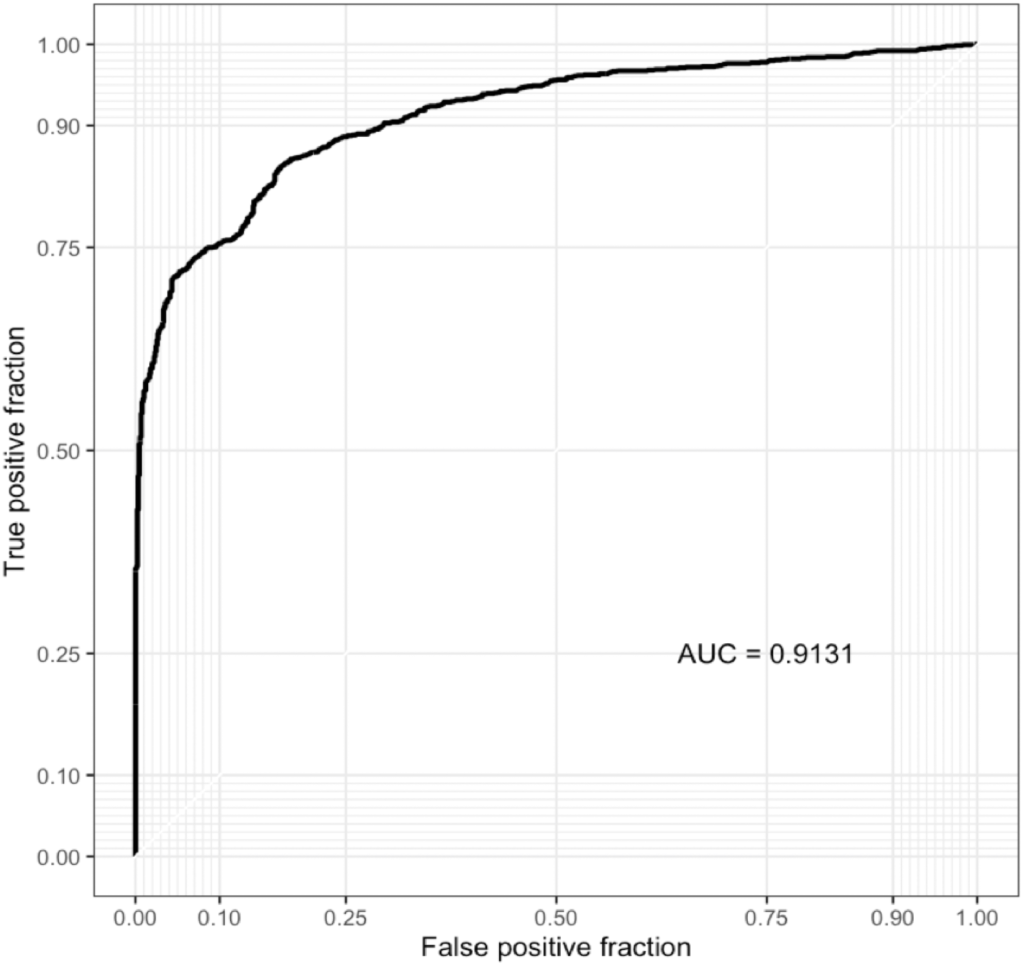
Biomarker ROC curve. ROC shows AUC and cross-validation prediction performance of the identified biomarker.

To summarize, with the help of *animalcules*, we explored and compared the microbial community difference between TB and control samples. Our analysis shows that the microbial community structure in the control group is more diverse and evenly distributed compared to the one in the TB group. Also, the TB group as well as control group each has a specific microbial composition that is shared within the group. Finally, we identified a subset of microbes that indicate its differentiating power between TB vs. control samples, which can be used as a new TB disease biomarker.

## Discussion

A fundamental characteristic of *animalcules* is its seamless interaction with the user through dynamic visualization tools. This design logic is rooted in the fact that researchers in microbiology must analyze their data at multiple levels (taxonomy) and multiple scales (normalization), thus data visualization and analysis become complicated without an organized analysis framework and workflow. *animalcules* solves this problem by providing a platform for interactively exploring large datasets, making it easier for users to identify patterns inherent in the dataset through appropriate analysis methods. Key analysis methods allow users to investigate differences in grouped relative abundance patterns between multiple sample groups in the phylum level, check the top abundant species in one specific sample group, or to check the individual sample-wise microbiome composition at different taxon levels. Patterns identified can be further tested through alpha/beta diversity statistical tests, differential abundance analysis, as well as biomarker identification. Furthermore, *animalcules* utilizes the MAE object, an efficient data structure for multi-omic sequencing data, which could be extended in the future to incorporate host sequencing assays, and enable compact methods for analyzing host-microbe interactions.

## Conclusion

In this report, we present *animalcules*, an open-source R package and Shiny application dedicated to microbiome analysis for both 16S rRNA and shotgun sequencing (metagenomics and metatranscriptomics) data. We incorporate leading and novel methods in an efficient framework for researchers to characterize and understand the microbial community structure in their data, leading to valuable insights into the connection between the microbial community and phenotypes of interest.

## Availability and requirements

Project name: *animalcules*

Project home page: https://compbiomed.github.io/animalcules-docs/

Operating system(s): Linux, OS X, Windows

Programming language: R

License: GNU GPLv3

## Declarations

### Acknowledgments

This research was supported by grants from the NIH R01 GM127430 and U01CA220413. The authors thank Lucas Schiffer (Boston University) as well as other members of the Johnson lab for helpful comments.

### Authors’ contributions

YZ, AF, SM contributed to the software development. YZ, AF, TF, SM and WEJ contributed to the writing and review of the manuscript before submission for publication. All authors read and approved the final manuscript.

### Availability of data and materials

animalcules is freely available on GitHub at https://github.com/compbiomed/animalcules or Bioconductor at https://bioconductor.org/packages/release/bioc/html/animalcules.html and is accompanied by comprehensive documentation and tutorials at https://compbiomed.github.io/animalcules-docs/

### Consent for publication

Not applicable.

### Ethics approval and consent to participate

Not applicable.

